# Investigating the correlation of muscle function tests and sarcomere organization in *C. elegans*

**DOI:** 10.1101/2021.04.13.439723

**Authors:** Leila Lesanpezeshki, Hiroshi Qadota, Masoud Norouzi Darabad, Karishma Kashyap, Carla M. R. Lacerda, Nathaniel J. Szewczyk, Guy M. Benian, Siva A. Vanapalli

**Affiliations:** Department of Chemical Engineering, Texas Tech University, Lubbock, TX, 79409, USA; Department of Pathology, Emory University, Atlanta, GA, 30322, USA; Department of Biological Sciences, Texas Tech University, Lubbock, TX, 79409, USA; MRC/Arthritis Research UK Centre for Musculoskeletal Ageing Research, University of Nottingham, United Kingdom & National Institute for Health Research Nottingham Biomedical Research Centre, Derby, DE22 3DT, UK; Ohio Musculoskeletal and Neurological Institute (OMNI) and Department of Biomedical Sciences, Ohio University, Athens, OH, 45701, USA

**Keywords:** Muscle physiology, burrowing assay, microfluidics, muscle genetics, sarcomere structure

## Abstract

**Background:** *Caenorhabditis elegans* has been widely used as a model to study muscle structure and function due to many genes having human homologs. Its body wall muscle is functionally and structurally similar to vertebrate skeletal muscle with conserved molecular pathways contributing to sarcomere structure, and muscle function. However, a systematic investigation of the relationship between muscle force and sarcomere organization is lacking. Here, we investigate the contribution of various sarcomere proteins and membrane attachment components to muscle structure and function to introduce *C. elegans* as a model organism to study the genetic basis of muscle strength.

**Methods:** We employ two recently developed assays that involve exertion of muscle forces to investigate the correlation of muscle function to sarcomere organization. We utilized a microfluidic pillar-based platform called NemaFlex that quantifies the maximum exertable force and a burrowing assay that challenges the animals to move in three dimensions under a chemical stimulus. We selected 20 mutants with known defects in various substructures of sarcomeres and compared the physiological function of muscle proteins required for force generation and transmission. We also characterized the degree of sarcomere disorganization using immunostaining approaches.

**Results:** We find that mutants with genetic defects in thin filaments, thick filaments and M-lines are generally weaker, and our assays are successful in detecting the functional changes in response to each sarcomere location tested. We find that the NemaFlex and burrowing assays are functionally distinct informing on different aspects of muscle physiology. Specifically, the burrowing assay has a larger bandwidth in phenotyping muscle mutants, because it could pick ten additional mutants impaired while exerting normal muscle force in NemaFlex. This enabled us to combine their readouts to develop an integrated muscle function score that was found to correlate with the score for muscle structure disorganization.

**Conclusions:** Our results highlight the suitability of NemaFlex and burrowing assays for evaluating muscle physiology of *C. elegans*. Using these approaches, we discuss the importance of the studied sarcomere proteins for muscle function and structure. The scoring methodology we have developed lays the foundation for investigating the contribution of conserved sarcomere proteins and membrane attachment components to human muscle function and strength.

## Background

Skeletal muscle, the major contractile tissue in animals, generates the force required for movement. The contractile force generation occurs by the sliding of two filaments composed primarily of the two most abundant muscle proteins - actin and myosin (1). Myosin head domains on the surface of thick filaments exert force by pulling oppositely oriented thin filaments primarily composed of actin inwards from either side, leading to shortening of the fundamental unit of contraction, the sarcomere. Since thousands of sarcomeres are connected end to end to produce myofibrils, and these myofibrils extend the length of the elongated skeletal muscle cell, this force is transmitted longitudinally within a muscle cell ultimately to the tendons and bones, to permit locomotion (1).

The contractile force generation and functional performance of the skeletal muscle can be significantly influenced by genetic defects in the contractile apparatus (2). Mutations such as in ACTN3 and ACE have been shown to improve muscle contractile performance in elite power athletes (3). Alternatively, some disease-causing mutations that impair human muscle function are well known such as in muscular dystrophies (4, 5), congenital myopathies (6), and cardiomyopathies (7). Since muscle strength is a useful predictor of all-cause mortality (8), there is a significant interest in uncovering the genetic basis for improved muscle mass (9) and strength (2) with age.

Understanding how genetic mutations influence muscle structure and function is best studied in model organisms (10). *Caenorhabditis elegans* with 95 body wall muscle cells required for whole animal locomotion is an excellent genetic model to study muscle organization, assembly, maintenance, and function (11, 12). Its sarcomere is made of dense body and M-line, in which the dense body is analogous to Z-line in humans. Thin filaments are anchored to dense bodies, while thick filaments are organized around M-lines. Actomyosin contractile forces are transmitted to the basement membrane and cuticle through the anchorage of both dense body and M-line that analogously serve the same function of costamere in vertebrate muscle, which are muscle-specific examples of integrin adhesion complexes (13). Unlike vertebrates, the contractile forces in *C. elegans* are mostly transmitted laterally to the cuticle resulting in bending and sinuous locomotion of the worm on an agar surface (14). In addition, integrin adhesion complexes exist at attachment plaques lying at the boundaries between adjacent muscle cells separated by basement membrane (15), and these also are likely to be involved in force transmission as their absence results in a locomotion defect (16).

The conserved molecular pathways contributing to sarcomere structure, and muscle function have been studied in *C. elegans* using a variety of methods. The animal’s transparent body along with immunostaining approaches and various microscopy techniques has helped to visualize the muscle structure (15, 17–21). Impairments in muscle function due to genetic defects have been studied by observation of gross phenotypes such as “Unc” (Uncoordinated) adults and “Pat” (Paralyzed arrest at 2-fold) embryonic lethals (13, 17, 22–24). In addition, thrashing and crawling assays have been used to assess muscle function in *C. elegans* (25, 26). Despite these advances, how the disorganization in muscle structure, due to genetic defects in the various components of sarcomeres and their membrane attachment structures, influences muscle function remains poorly understood.

Recently, new methods have been developed in *C. elegans* that focus on quantitative characterization of muscle function with the capacity to detect more subtle phenotypes in muscle mutants. A bending amplitude assay has been used to identify the locomotion defects in muscle focal adhesion mutants (27). Optogenetic-stimulation has also been used to measure muscle contraction/relaxation rate constants by monitoring body area changes (28). Assessing the animals push and pull forces using atomic force microscopy (29) and pillar deflection measurements in microfluidic devices (30–32) have introduced sophisticated means to calculate the muscle strength of *C. elegans*. Additionally, characterizing the burrowing performance of *C. elegans* where animals move in 3D have enabled assessment of neuromuscular health (33, 34).

Studies have begun to employ these new methods to address the relationship between muscle structure and function in *C. elegans*. Disorganization of actin filaments could support that glucose-treated wild-type animals have reduced thrashing force (32), and electron microscopy of body wall muscle cross-section could justify that the low muscle force is due to disorganization in dense bodies and M-lines (29). However, studies lack a systematic investigation of the relationship between muscle force and structural disorganization which is introduced due to a variety of genetic defects in sarcomere and membrane attachment components. Here, we have utilized two recently developed assays in our laboratory that report on muscle function. The first assay involves measurement of muscle strength using a micropillar-deflection based system called NemaFlex (31) and the second assay involves stimulating animals to burrow in three dimensions in a resistive gel medium (34). We implemented these assays on a set of 20 muscle mutants with genetic defects in various sarcomere components. We also characterized the degree of sarcomere disorganization in these mutants and tested the ability of these assays to correlate muscle function to muscle structure. Our results highlight the suitability of our assays and scoring approaches for evaluating *C. elegans* as a genetic model for muscle strength.

## Methods

### Worm culture

*Caenorhabditis elegans* animals were cultured at 20 °C on standard nematode growth medium (NGM) on 60 mm petri plates and never allowed to starve. The NGM plates were seeded with *Escherichia coli* OP50 bacteria overnight. Age synchronization was done by transferring 20-25 gravid animals to seeded plates and letting them lay eggs for ~3 hours. The gravid adults were then removed, leaving age synchronized eggs to hatch and develop at 20 °C until they reach day 1 of adulthood for all the experiments. Day 0 of adulthood is when the age synchronized animals started to lay eggs.

*unc-22(sf21), unc-94(sf20)*, and *unc-98(sf19)* were generated in and available in the Benian lab. Wild-type, N2(Bristol), and the following mutants were obtained from *Caenorhabditis* Genetics Center: *dyc-1(cx32), pfn-3(tm1362), uig-1(ok884), atn-1(ok84), zyx-1(gk190), unc-95(ok893), tln-1(e259), unc-82(e1220), unc-89(e1460), pkn-1(ok1673), alp-1(tm1137), dim-1(ra102), unc-22(e66), unc-54(s95), unc-54(s74), lev-11(x12), and unc-60(r398)*. Nearly all of these mutants had been multiply outcrossed to wild type.

### Pluronic gel-based burrowing assays

Burrowing assays were conducted as previously described (34). Briefly, 26 % w/w Pluronic F-127 (Sigma-Aldrich) solution was prepared and stored at 4 °C prior to the experiment to prevent gelation. A minimum number of 30 animals were transferred into the bottom of a Corning™ Falcon™ Polystyrene 12-well plate at 20 ± 1 °C, either by handpicking them into 20-30 μL Pluronic solution or in 10 μL of worm solution in water, which was then combined with 500 μL of PF-127 to make a base gel layer. Then, the Pluronic layer was cast on top to the thickness of 0.7 cm, followed by 20 μL of 100 mg/mL *E. coli* solution in liquid NGM as an attractant (t=0 min). The number of animals burrowed to the surface was scored every 15 minutes for a total duration of 2 hours. Three replicates per strain were conducted. To compare mutants burrowing performance, two-way ANOVA was used in GraphPad Prism software.

### *C. elegans* muscle strength measurements using NemaFlex

The muscle strength of *C. elegans* strains was measured using the NemaFlex technique as previously described (31), based on the deflection of soft micropillars as the animals are crawling through the pillar arena. Polydimethylsiloxane (PDMS) devices were poured (Sylgard 184 part A (base) and part B (curing agent) 10:1 by weight; Dow Corning) over the mold by curing for 2.5 hours at 70 °C. The micropillar devices had pillars arranged in a square lattice with a pillar diameter of 44 μm and a height of 87 μm. The gap between the pillars is 71 μm.

Synchronized day 1 adults were loaded individually in each food-free chamber (35), followed by a 1-minute video collected for each animal at 20 ± 1 °C. Imaging was performed in brightfield using a Nikon Ti-E microscope with a 4x objective and Andor Zyla sCMOS 5.5 camera at 5 frames per second and a pixel resolution of 1.63 μm per pixel. Movies were processed and analyzed for strength values using our in-house-built image processing software (MATLAB, R2016a) (Available at https://github.com/VanapalliLabs/NemaFlex). Animal strength was calculated by identifying the pillar with the maximum deflection in each frame to estimate maximal force exerted. We bin these maximal forces and define the animal muscular strength as *f*_95_, which corresponds to the 95^th^ percentile of these maximal forces. We normalized *f*_95_ by the cube of animal body diameter to account for differences in animal body diameter (31). The muscle strength of mutants was compared with WT animals by calculating the muscular strength ratio and denoting it as the fold change in muscle strength. Statistical analysis was performed using Wilcoxon rank-sum test in MATLAB.

### Immunostaining of body-wall muscle

Adult nematodes were fixed and immunostained as described previously (36, 37). The following primary antibodies were used: anti–ATN-1 (Mouse monoclonal MH35 (19); kindly provided by Dr. Pamela Hoppe, Western Michigan University) and anti–myosin heavy chain A (MHC A; mouse monoclonal 5-6 (38); purchased from the University of Iowa Hybridoma Bank) were used at 1:200 dilution, anti–UNC-89 (Rabbit polyclonal EU30) (39)) and anti–UNC-95 (Rabbit polyclonal Benian-13 (40)) were used at 1:100 dilution. Secondary antibodies included anti-rabbit Alexa 488 (Invitrogen), and anti-mouse Alexa 594 (Invitrogen) both diluted at 1:200. Rhodamine-phalloidin staining of thin filaments was carried out as described (41). Images were captured at room temperature with a Zeiss confocal system (LSM510) equipped with an Axiovert 100M microscope and an Apochromat 63x/1.4 numerical aperture oil immersion objective in 2.5x zoom mode. The color balances of the images were adjusted using Adobe Photoshop (Adobe, San Jose, CA).

### Muscle disorganization score calculation

The muscle disorganization score was generated from assessing the immunostaining results of this study (not including others) in which the same methods and antibodies for immunostaining were used. In addition, the scoring was performed independently by two investigators, and the images presented were representative examples from 10 worms for each strain. The discrepancy between two different observers on sarcomere disorganization have been reported to have the coefficient variation of less than 15% (42).

## Results

### Selection of muscle mutants and prior assessment of muscle function

To investigate the relationship between muscle structure and function in *C. elegans*, we selected 18 muscle proteins (Figure 1). This selection was based on (i) proteins that were located on different structural components of the sarcomere, (ii) mutants that have been previously characterized in terms of locomotion (Table 1), (iii) whether these proteins are involved in generating muscle forces versus transmitting them. With respect to proteins involved in force generation, we selected 2 proteins associated with thick filaments (UNC-22 and UNC-54); and three proteins associated with thin filaments (LEV-11, UNC-60, and UNC-94). With respect to force transmission, we selected 6 proteins localized in dense bodies (UIG-1, DYC-1, PFN-3, ATN-1, ALP-1, and DIM-1) and 3 proteins localized in M-lines (UNC-82, UNC-89, UNC-98). We also selected 4 proteins that are localized in both dense bodies and M-lines (ZYX-1, TLN-1, UNC-95, PKN-1). Although our selection is limited, targeting various structural units of the sarcomere provides an initial assessment of the role of these proteins in muscle function.

**Figure 1:**
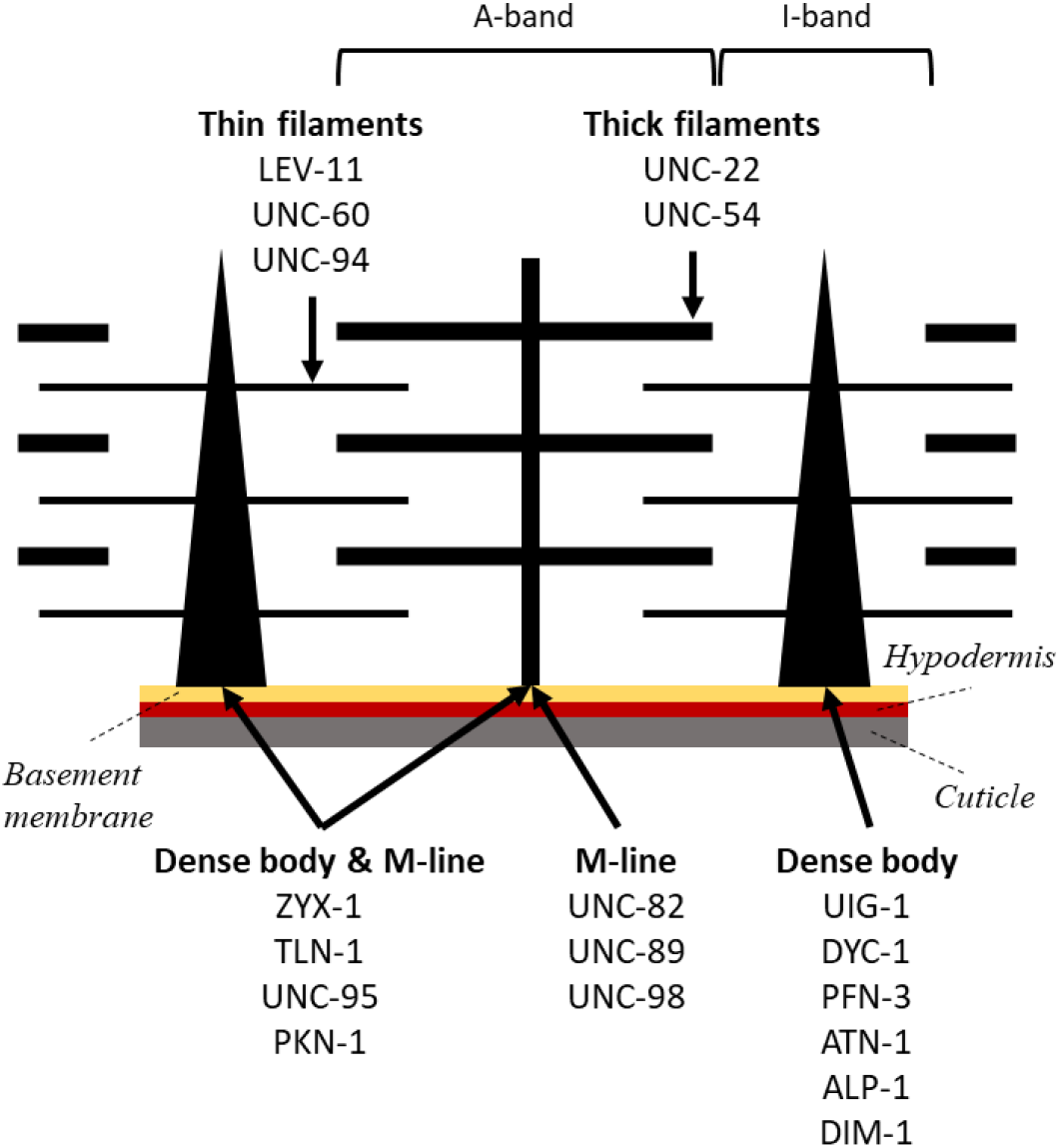
The *C. elegans* sarcomere highlighting the various structural components of the sarcomere. The muscle mutants were selected based on the genes encoding proteins localized to the 5 different structural sites of the muscle: thin filaments, thick filaments, M-line, dense body, and both dense body and M-line.

**Table 1.** Summary of the mutants studied, and their human homologs, protein location and reported behavioral phenotypes. *The crawling assays were conducted under stimulated conditions. ** http://www.wormbase.org, release WS274

| Mutant | Type of mutation | Human homolog | Protein location | Phenotype |  |  |
| --- | --- | --- | --- | --- | --- | --- |
|  |  |  |  | Body bending | Swimming | crawling |
| *uig-1(ok884)* | deletion, null | PLEKHG1, PLEKHG3 | dense bodies | normal (27) | normal (28) | fast*(28) |
| *pfn-3(tm1362)* | deletion, null | profilin | dense bodies | defective (27) | slightly slow (43) | -- |
| *atn-1(ok84)* | deletion, null | α-actinin | dense bodies | defective (27, 44) | slow (28) | normal*(28) |
| *alp-1(tm1137)* | deletion, null | cypher/ZASP/ oracle | dense bodies | defective (27) | normal (45) | -- |
| *dyc-1(cx32)* | unknown | capon | close to dense bodies | overbent (46) | -- | fast (46) |
| *dim-1(ra102)* | Splice site; loss of function | -- | around and between dense bodies | defective (27) | slow (28) | fast*(28) |
| *zyx-1(gk190)* | deletion, null | Zyxin | dense bodies & M-lines | defective (27) | -- | -- |
| *tln-1(e259)*<br>previously,<br>*unc-35* | unknown | Talin | dense bodies & M-lines | overbent** | -- | slow (47) |
| *unc-95(su33)* | nonsense, null | -- | dense bodies & M-lines | defective (27) | -- | slow (22) |
| *pkn-1(ok1673)* | deletion, null | protein kinase N | dense bodies & M-lines | -- | slow (48) | -- |
| *unc-82(e1220)* | missense | ARK5 & SNARK protein kinases | M-lines | -- | -- | slow (47) |
| *unc-89(e1460)* | nonsense in alternatively spliced exon; hypomorph | obscurin | M-lines | -- | slow (49) | slow (49) |
| *unc-98(sf19)* | splice site; loss of function | -- | M-lines | normal (27) | slow (28, 50) | normal (28) |
| *unc-22(sf21)* | missense | titin family member | thick filaments | -- | fast (51) | fast (51) |
| *unc-22(e66)* | 2 bp deletion; frame-shift to nonsense; loss of function | titin family member | thick filaments | -- | slow (28, 51) | slow (51) |
| *unc-54(s95)* | missense | myosin heavy chain | thick filaments | -- | -- | slow (52) |
| *unc-54(s74)* | missense | myosin heavy chain | thick filaments | -- | slow (28) | slow (28, 52) |
| *lev-11(x12)* | Missense<br>note: null is Pat embryonic lethal | tropomyosin | thin filaments | -- | normal (28) | fast*(28) |
| *unc-60(r398)* | nonsense near C-terminus; hypomorph | ADF/cofilin | thin filaments | -- | slow (28, 53) | slow*(28) |
| *unc-94(sf20)* | nonsense; null | tropomodulin | thin filaments | -- | slow (54) | -- |

Table 1 shows the list of genes encoding the selected proteins, the particular mutant alleles chosen, indication of the proteins’ human homologs, location in the sarcomere, and phenotypic information from locomotory assays reported in the literature. For two genes, *unc-54*, which encodes the major myosin heavy chain of body wall muscle thick filaments, and *unc-60*, which encodes ADF/cofilin, we chose to study mild alleles, as the null state for either results in severe paralysis.

The compiled phenotypes include the maximum bending amplitude from a bending assay (27), and the movement rate evaluated from standard swimming and crawling assays. Examining the data, we find that the mutants belonging to M-lines and dense bodies & M-lines showed slow crawling or swimming movements (Table 1), except for *unc-98(sf19)* which showed normal crawling but slower movement in swimming. On the contrary, mixed phenotypes were observed for dense bodies, thick filaments, and thin filaments mutants. Some mutants did not show any apparent defects in crawling or swimming assays and some were even better than wild-type (WT) animals. Mutants *uig-1, dyc-1, dim-1*, and *lev-11* were faster in crawling only and *unc-22(sf21)* was faster in both assays. On the other hand, bending assays identified six of the mutants to be defective in bending (*zyx-1, unc-95, pfn-3, atn-1, alp-1, dim-1*) and two were normal (*unc-98, uig-1*). Interestingly, both *tln-1* and *dyc-1* showed exaggerated body bends; however, their movement rates were different with *tln-1* crawling slowly, and *dyc-1* crawling faster.

### Genetic defects in sarcomeres can reduce *C. elegans* muscle strength and burrowing prowess

Due to the localization and known function of these proteins in the sarcomere, it is expected that their corresponding loss-of-function mutants would show muscle weakness than WT. However, none of the locomotory assays listed in Table 1 are suitable for measuring muscle strength. Previously, the NemaFlex assay was used to show muscle weakness in *unc-17(e245), unc-52(e669), unc-112(r367ts)* and multiple alleles of *unc-89* (31, 49). Here, we have utilized the NemaFlex platform to measure strength of many more muscle mutants listed in Table 1. Since the body diameter of some of the mutants was significantly different compared to WT (see SI Table S1), we normalized the muscle strength by the cube of the body diameter. This normalization was previously shown to account for the influence of body diameter reasonably well (31) and it also works well for the muscle mutants studied here (see SI Figure S1).

In Figure 2B, we show the normalized strength data for the tested mutants. We find that the mutants with defects in thin filaments, thick filaments and M-lines were generally weaker than WT except *lev-11(x12), unc-22(sf21), unc-54(s74)* and *unc-98(sf19)*. Animals with genetic impairments in both dense body and M-line did not show a significant difference compared to WT except *unc-95(su33)*. Finally, four out of the six mutants with genetic defects in dense bodies did not show a difference while the other two were weaker than WT. In addition, we find that out of the 20 tested mutants, *unc-95* was the weakest.

**Figure 2:**
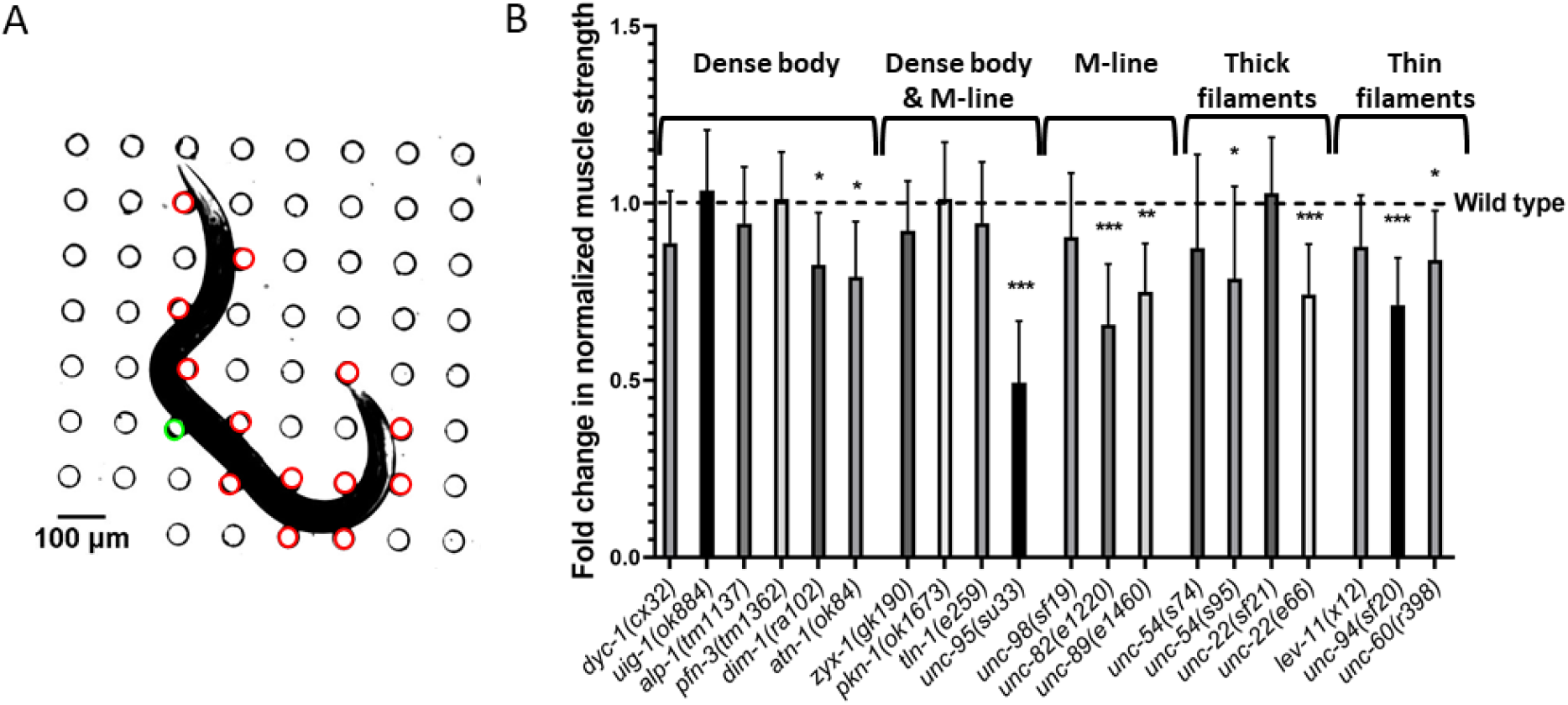
NemaFlex strength measurements on muscle mutants. (A) An animal is crawling in a NemaFlex chamber and deflecting the pillars. The red pillars are in contact with the worm body and undergoing deflection due to muscle forces. The green pillar has the maximal deflection or maximal force exerted by the animal. (B) The fold change in normalized muscle strength of mutants compared to wild-type animals. See Materials and Methods section for estimation of muscle strength from maximal pillar forces and normalization. Error bars represent standard error of the mean. Significance levels are assessed with Wilcoxon rank-sum test, *P ≤ 0.05, **P ≤ 0.01, ***P ≤ 0.001. Sample sizes were as following: WT had an average sample size (N) of 30. N= 36, 30, 26, 25, 31, 29 for *dyc-1, uig-1, pfn-3, dim-1, atn-1*. N= 20, 25, 34, 20 for *zyx-1, pkn-1, tln-1, unc-95*. N= 27, 29, 31 for *unc-98, unc-82, unc-89*.N= 36, 34, 26, 31 for *unc-54(s74), unc-54(s95), unc-22(sf21), unc-22(e66)*. N= 27, 23, 27 for *lev-11, unc-94, unc-60*.

The results of Figure 2 show that 9 out of 20 tested muscle genes contribute to muscle strength with 6 out of the 9 belonging to genetic defects in M-lines, thick and thin filaments. It appears that genetic defects in these sarcomere components might lead to a more severe loss in muscle strength, thus the assay can detect the physiological consequences of genetic defects in each of these structures. It is interesting to note that the null mutations of 3 components of dense body (*uig-1, pfn-3, alp-1*) each do not result in severe defects compared to mild mutations of the other structures. Thus it appears that dense body is more resilient to loss of individual components with respect to strength production.

Complementing the NemaFlex assay, we also tested mutants in the burrowing environment where the muscles of *C. elegans* might be challenged differently due to the three-dimensional movement (55, 56). In the burrowing assay, mutants are loaded at the bottom of well-plate and then stimulated to burrow through the Pluronic gel towards an attractant on the top (Figure 3A). The percentage of the animals that could reach the top are counted at 15-minute intervals for a total duration of 2 hours (34).

**Figure 3:**
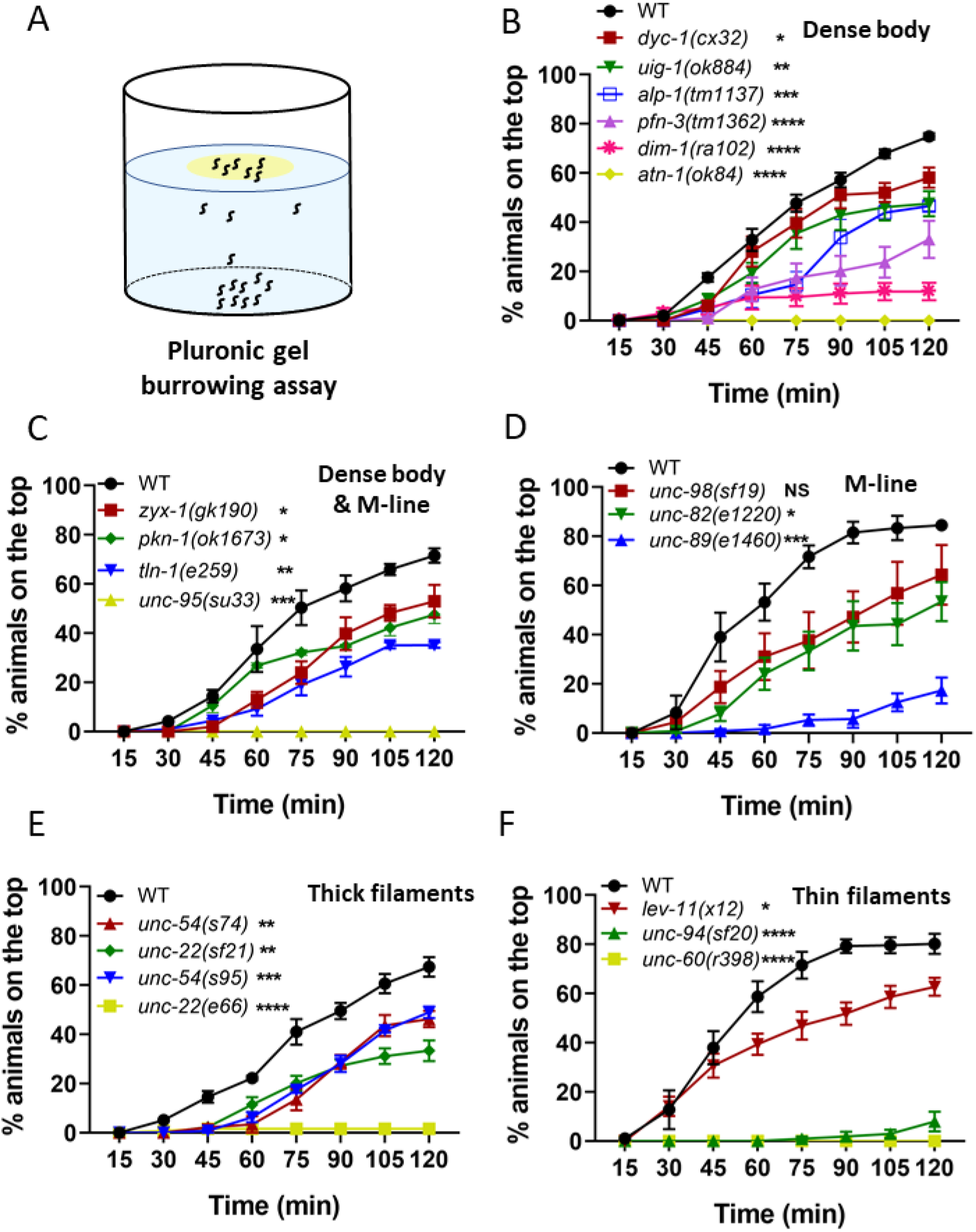
Burrowing performance of muscle mutants. (A) In a Pluronic gel-based burrowing assay, the animals are loaded on the bottom of a well-plate and stimulated to burrow toward an attractant (in yellow) on the top. (B-D) The burrowing performance was assessed by determining the percentage of the animals on the top for 2 hours. The mutants were with genetic defects in the sarcomere genes located on (B) dense body, (C) dense body and M-line, (D) M-line, (E) thick filaments, and (F) thin filaments. Sample sizes were as following: (B) N = 30-44 for WT, 31-40 for *dyc-1*, 31-43 for *uig-1*, 45-54 for *alp-1*, 29-39 for *pfn-3*, 41-48 for *dim-1*, 29-40 for *atn-1*. (C) N = 32-41 for WT, 30-35 for *zyx-1*, 35-40 for *tln-1*, 33-36 for *unc-95*. (D) N = 33-41 for WT, 26-39 for *unc-98*, 41-45 for *unc-82*, 28-41 for *unc-89*, (E) N= 45-49 for WT, 39-56 for *unc-54(s74)*, 44-50 for *unc-22(sf21)*, 41-52 for *unc-54(s95)*, 39-58 for *unc-22(e66)*, (F) N = 40-48 for WT, 31-35 for *lev-11*, 30-35 for *unc-94*, 34-43 for *unc-60*. Minimum of 3 replicates were conducted per strain. Error bars are standard error of the mean. Significance levels are assessed with two-way ANOVA.

The burrowing assay was used previously to show that 4 dense body, and 3 dense body and M-line protein mutants burrow less effectively than WT (34), which led us to evaluate additional muscle mutants with this assay. As shown in Figure 3B-F we find that most of the muscle mutants tested could not burrow as effectively as wild-type, except *unc-98(sf19)*. There was no obvious correlation between the burrowing performance and the structural component where defects were present. Interestingly, none of *atn-1(ok84), unc-95(su33)* and *unc-60(r398)* animals could reach the attractant on top, making them amongst the worst burrowers, followed by *unc-22(e66)* and *unc-94(sf20)* where only 2% and 7% of the animals reached the attractant (Figure 3B).

We have previously shown that this burrowing assay is not merely a chemotaxis assay and in fact, involves exertion of muscles since the burrowing performance varied with gel stiffness and height (34). Also, the mutants that are not defective in 2D chemotaxis (Fig. 6c in reference 34) showed deficiency in burrowing indicating that the muscle actuate differently during the 3D locomotion providing more information on muscle function than 2D chemotaxis assays. This is also confirmed in this study by testing Unc mutants that are known to have uncoordinated movements; however, in a 2D chemotaxis assay, none of *unc-22* and *unc-54* mutants had any significant impairment to control their movement toward the bacteria in 2D (Figure S2), yet all are defective in burrowing assay.

Overall, both the NemaFlex and burrowing assays inform on the muscle function of *C. elegans* and highlight that genetic impairments in the muscle can lead to loss of muscle performance.

### Burrowing and NemaFlex strength measures are functionally distinct

To develop an integral measure of muscle function that combines readouts from NemaFlex and burrowing assay, we first considered whether readouts from these two assays report on distinct aspects of muscle function. The locomotory forces generated by *C. elegans* during two-dimensional motion through a mechanical environment consisting of deformable micropillars is expected to be different than those generated during three-dimensional maneuvers made during burrowing, suggesting that these assays might report on unique aspects of muscle function. Indeed, we showed recently that measures extracted from NemaFlex and burrowing for WT animals do not correlate with one another (56). Here, we sought to address the existence or lack of this correlation for the muscle mutants. In addition, we were interested to check if there is data clustering based on the known location of the protein in the sarcomere that is affected by the mutation.

We calculated Z-scores 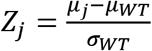 with respect to the wild-type population for each of the functional measures extracted from the NemaFlex and burrowing assay. Here *μ* is the mean measured value, which in our case is either the normalized muscle strength (from the NemaFlex assay) or the percent animals reaching the surface at the 2 hr time point (from the burrowing assay), and the subscript *j* denotes a given mutant strain. Likewise, *μ_WT_* and *σ_WT_* represent the mean and the standard error of the measured values for WT animals. Thus, a Z-score of 0 indicates that the mutant’s muscle function measures were identical to WT.

In Figure 4, we show the Z-scores from the burrowing and NemaFlex assays for all the tested mutants. In the previous section, we showed *unc-95, atn-1, unc-60, unc-22(e66)*, and *unc-94* were the worst burrowers. Consequently, all of these mutants are found on the far left of Figure 4; however, their NemaFlex strength measurements vary from *Z* ≈ −3 for *unc-60*, to the weakest mutant in both NemaFlex and burrowing assays, *unc-95*, with *Z* ≈ −8. On the other hand, *dyc-1, uig-1, alp-1, pfn-3, zyx-1, pkn-1, tln-1, unc-98, unc-54(s74), unc-22(sf21), lev-11* that were not significantly different than WT in strength measurement lie toward the top right side of Figure 4 with a large range of burrowing Z-score between −19 for *unc-22(sf21)* to −4.8 for *unc-98*. This suggests that the muscle function impairments are more pronounced in burrowing assay. Overall, this data shows that these two assays are modestly correlated (Spearman correlation coefficient is 0.42) which is expected since both involve actuation of muscles, but given that they are not completely correlated also suggests that they can inform on different aspects of muscle physiology. We also could not find any evidence of clustering based on the locations of the proteins affected.

**Figure 4:**
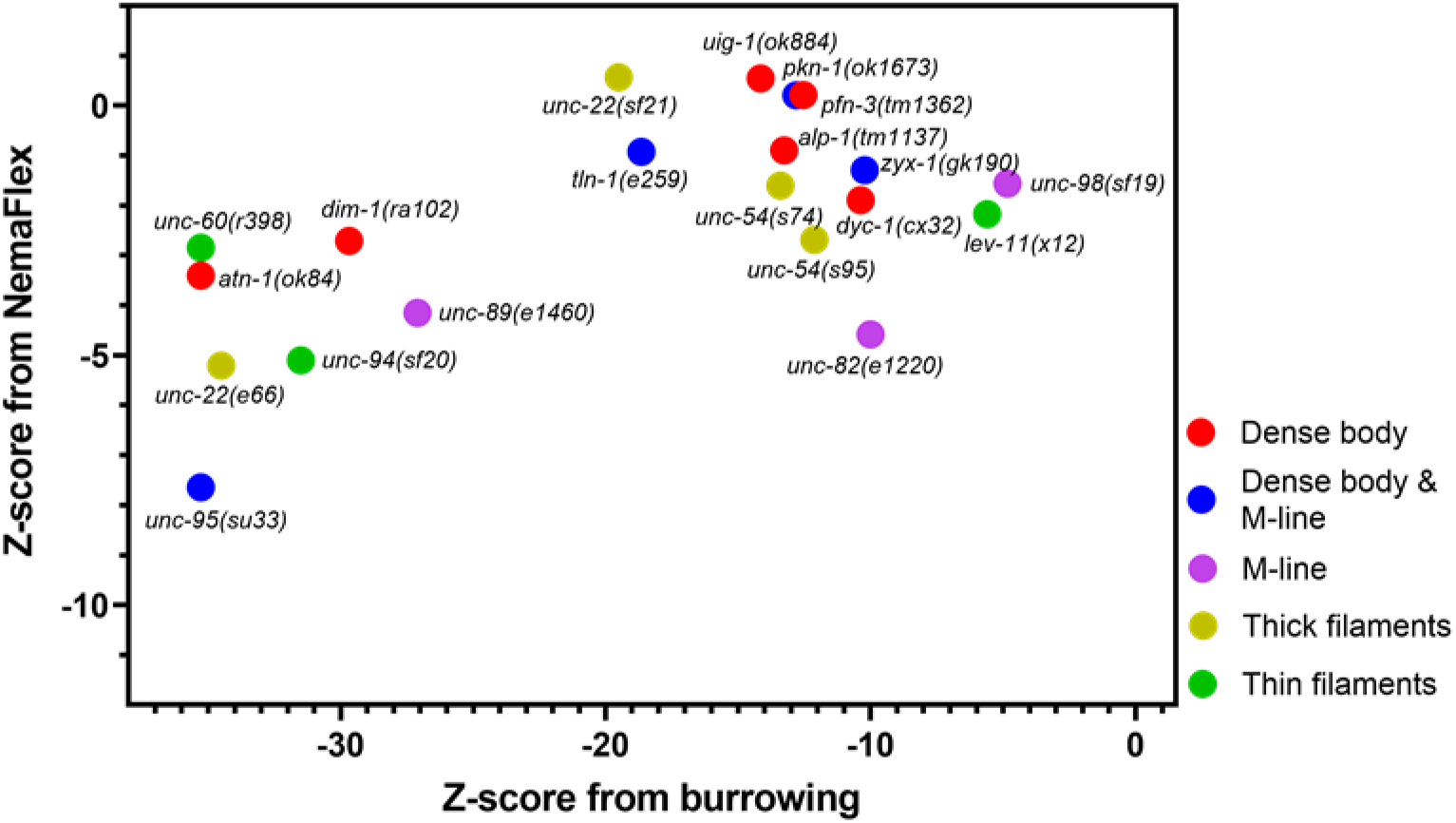
Burrowing assay and NemaFlex strength measurement are not correlated strongly. Z-score of 0 represents being identical to WT. The mutants are color coded based on the protein location in the sarcomere.

### A scoring system for structural disorganization of *C. elegans* muscle

Structural disorganization in the sarcomeres can adversely affect the contraction-relaxation cycle consequently interfering with muscle function. As NemaFlex and burrowing assays engage *C. elegans* in actuating its muscles, we speculated that the mutants with a low Z-score in at least one of the assays are likely to exhibit some level of disorganization in the sarcomere structure. We therefore performed immunostaining to visualize the structural disorganization in the muscle contractile apparatus of mutants and developed a scoring system to rank the level of disorganization.

Dense body, M-line, A-band, and I-band are the main substructures of the sarcomere, the fundamental unit of muscle contraction. Thus, we evaluated sarcomeric organization at these sites by immunostaining with antibodies to the following proteins: MHCA (myosin heavy chain A) that is located in the middle of A-bands which represent assembled and organized myosin thick filaments, UNC-95 that is located at the bases of dense bodies and M-lines, UNC-89 that is located throughout the depth of the M-lines, and ATN-1 that is located in the major but not basal portion of dense bodies. Phalloidin was also used to stain I-bands where actin thin filaments are located. We also surveyed the literature and collected images of the aforementioned muscle structures of the same allele used in this study. Table 2 shows the muscle structure data for 20 mutants with 69 data points obtained from this study, and 36 collected from literature. The immunostained images for the mutants obtained in this study are shown in SI Figure S3.

**Table 2.**
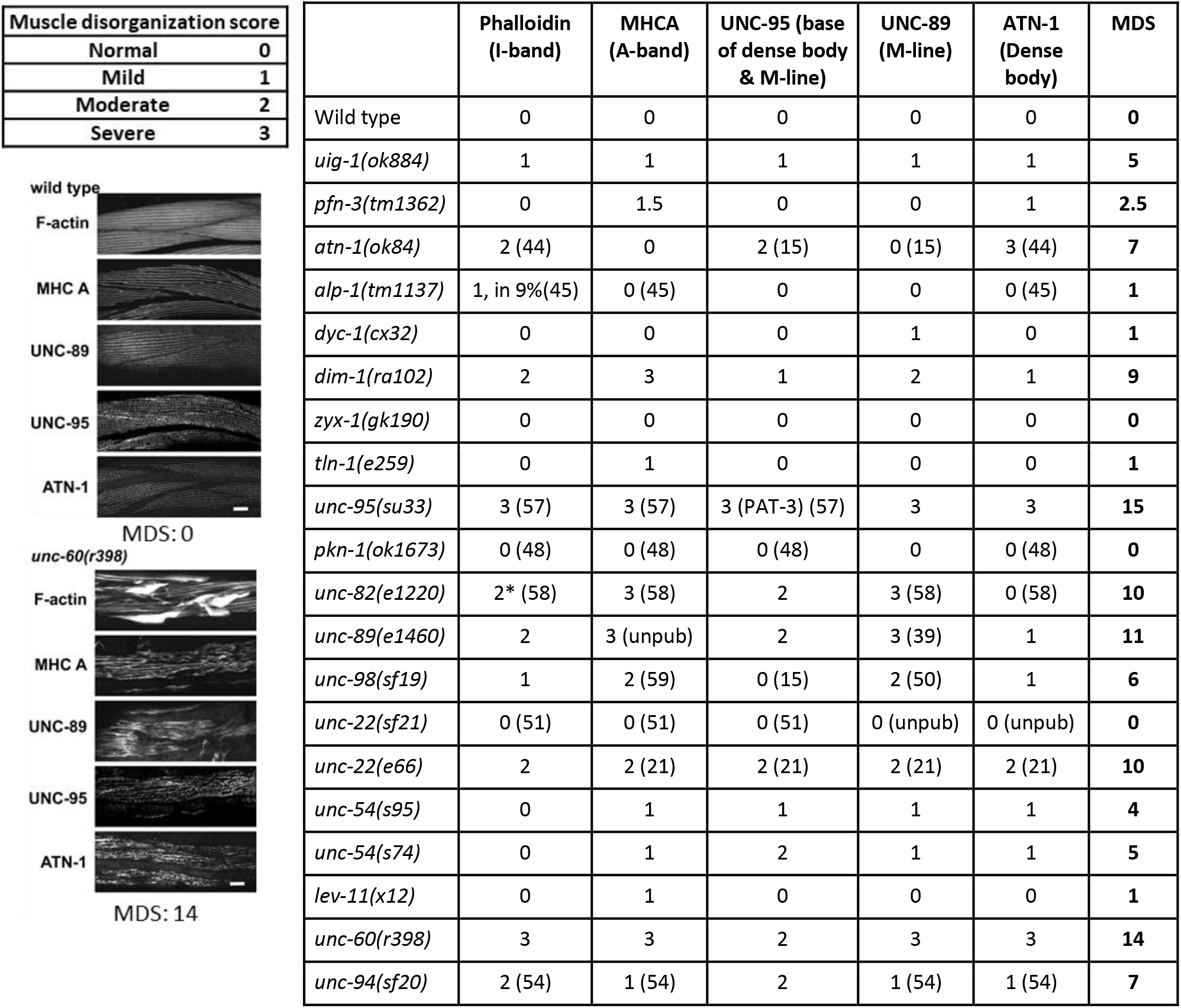
Muscle Disorganization Score (MDS). Each of the structural sites were given a score from 0 to 3, based on the severity of disorganization. MDS for a mutant is the sum of these scores on each site. Left panel refers to WT with normal muscle structure and MDS = 0, while this score is 14 for *unc-60(r398)* due to the severity of its muscle disorganization. *Immunostaining results reported in (Hoppe et al. 2010) were obtained on a different allele, *unc-82(e1323)*, which is a premature stop codon and is likely a null allele.

The mutants were evaluated based on the level of structural disorganization of each of the stained structures and were assigned a score from 0 to 3 for normal to the most severe disorganization (Table 2). Muscle Disorganization Score (MDS) is defined as the sum of the disorganization score of each individual structure. A mutant can acquire a value ranging from 0 for an organized structure to a maximum of 15 for severely disorganized muscle structure. The left panel of Table 2 illustrates two examples on how MDS is calculated: WT animals or the control group received an MDS = 0, as none of the structures studied were disorganized, and *unc-60(r398)* was given MDS = 14 because of the severe structural disorganization observed on I-bands, A-bands, and the depth of M-lines and dense bodies, along with the moderate disorganization on the base of dense bodies and M-lines.

With respect to mutants, we did not observe any disorganization in *zyx-1, pkn-1, unc-22(sf21)* body wall muscle structure (Table 2). Interestingly, none of these mutants were weaker in NemaFlex strength measurements, yet they were all burrowing impaired. Likewise, the mutants with low Z-scores on both NemaFlex and burrowing shown on the bottom left corner of Figure 4, had structurally disorganized muscles; *i.e*. MDS of *unc-95* and *unc-22(e66)* were 15 and 10, respectively. In the next section, we use this scoring system to understand how muscle structure and muscle function are related to each other.

### Correlating muscle function with muscle structure disorganization

To understand how structural organization of the myofilament lattice can contribute to muscle function, we sought a functional readout characterizing muscle physiology. Since NemaFlex and burrowing Z-score values were not in the same range, we normalized them with respect to the absolute maximum so that the normalized Z-scores 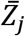 lie in the range 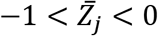. In Figure 5A and Figure 5B, we plot the normalized NemaFlex and burrowing scores as a function of the muscle disorganization score. In general, both the normalized NemaFlex and burrowing scores decrease with increase in the score of muscle structure disorganization. Principal component analysis (PCA) suggested that the summation of the normalized NemaFlex and burrowing Z-scores with almost the same weight, can result in a new latent variable, which we refer to as muscle function score that could explain 80% of data variation (See SI Note S1). Figure 5C shows that the correlation using this new muscle function is improved compared to each individual physiological Z-score. Interestingly, all data points fall inside the 95% prediction bound of the fitted line. Overall, these results suggest that the lower the muscle function score is, the greater is the degree of sarcomeric disorganization, indicating these two muscle physiology assays together can improve the chance of successfully detecting any structural disorganizations in the animal’s body wall muscle.

**Figure 5:**
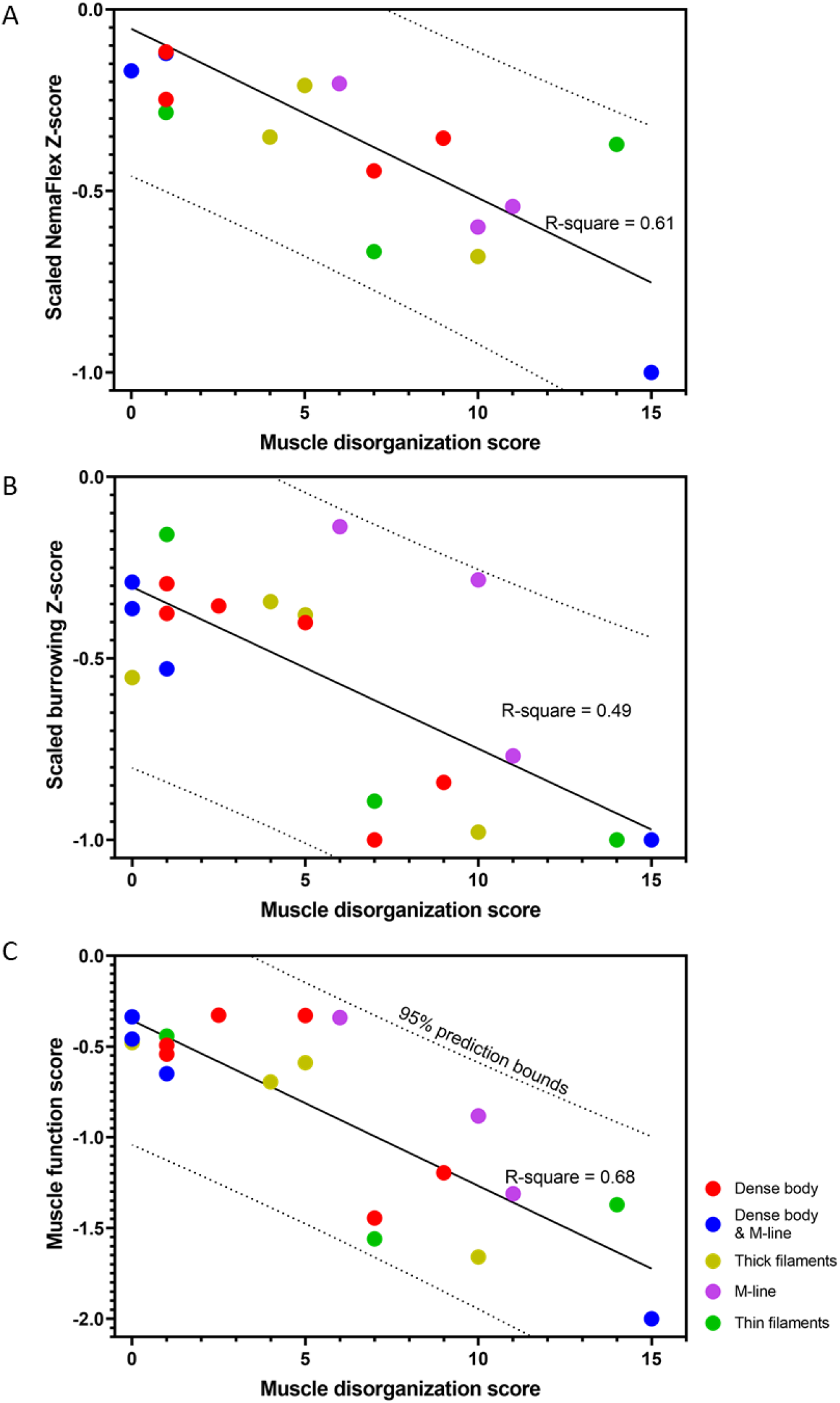
Muscle function score is correlated with muscle disorganization score. (A,B) NemaFlex and Burrowing Z-score values show the R-square of 0.61 and 0.49 respectively from linear regression. (C) Muscle function score is the summation of scaled Z-scores from NemaFlex and burrowing assays. The black line shows the linear regression fit with R-square of 0.68 and the 95% prediction bounds are shown as dotted lines.

Additionally, Figure 5C shows that all the M-line and thick filaments mutants are mildly disorganized in their muscle structure with mixed physiological functions. The muscle structure of dense body mutants varies from nearly normal values to mild disorganization and also normal to mild physiological function. Most of the dense body & M-line mutants have almost well-organized muscle structures with normal physiological performance, except for *unc-95* that was both structurally and physiologically severely impaired. For mutants with defects in thin filaments, muscle structural disorganization and muscle function vary from mild to severe.

## Discussion

### Novel assays advance studies of muscle function in *C. elegans*

Our results demonstrate the effectiveness of two novel assays - NemaFlex and burrowing - in detecting functional deficiencies in *C. elegans* muscle mutants. Furthermore, our results suggest that each of the assays informs on different aspects of muscle physiology. Thus, assays in which *C. elegans* response is measured in 2D and 3D environments can provide unique information on muscle function.

Both the NemaFlex and burrowing assays showed 9 mutants out of 20 being defective. This includes *dim-1* and *atn-1* with defects in dense body, *unc-95* with a defect in dense body and M-line, *unc-82* and *unc-89* with defects in M-line, *unc-54(s95)* and *unc-22(e66)* with defects in thick filaments, and *unc-94* and *unc-60* with defects in thin filaments (Figures 2 and 3). Apart from these mutants, there were 10 other mutants that were only burrowing impaired. This shows that the burrowing assay has a larger bandwidth in phenotyping muscle mutants with subtle genetic defects which are not detected by NemaFlex.

Comparing the outcomes from standard locomotory assays of swimming or crawling, we observe that out of the 20 muscle mutants, except four mutants *uig-1, dyc-1, unc-22(sf21)* and *lev-11(x12)*, the rest are defective in either crawling or swimming (Table 1). 9 out of the 20 muscle mutants that were found to be defective in both NemaFlex and burrowing were also found to be defective in swimming or crawling. Interestingly, among the four mutants showing a fast phenotype in either the crawling or swimming assay, *uig-1, dyc-1, unc-22(sf21)* and *lev-11(x12)* were significantly burrowing impaired, yet none of them were weaker in NemaFlex muscle strength assessment.

Overall, comparing the results from NemaFlex and burrowing assays to the traditional swimming and crawling assays suggests that the common impairments among all these assays are mostly associated with the mutations of the related proteins found on M-line in 2 mutants, dense body and M-line in 1 mutant, thick filaments and thin filaments each in 2 mutants. Mixed phenotypes were identified in dense body mutants, as there was no concurrence in *atn-1* and *dim-1* swimming and crawling assays, yet both were impaired in NemaFlex and burrowing.

### Correlating muscle function to muscle structure

We investigated mutants with genetic defects in their muscle structure with the aim of relating structural disorganization in sarcomeres to muscle function. We built a scoring system to evaluate the muscle structural disorganization by immunostaining key sarcomeric structural components including dense body, M-line, A-band, and I-band. We also developed a muscle function score by combining the outputs of both NemaFlex and burrowing assays to result in a score between −2 for the most impaired animals and 0 for the animals that were physiologically similar to WT.

Our results indicate that the muscle function score is negatively correlated with muscle disorganization score (Figure 5C). *Unc-95, unc-60, unc-89, unc-82*, and *unc-22(e66)* were the mutants with the highest level of disorganized body wall muscles with muscle disorganization score of 15, 14, 11, 10, 10, respectively. These mutants can be found on the lower right of Figure 5C, correlated with low muscle function score justifying their impaired muscle function. Interestingly, *atn-1* and *unc-94* were found to be physiologically impaired (score of ~-1.5 out of the minimum −2), yet their muscle structure was only mildly disorganized.

Mutants *zyx-1, pkn-1, unc-22(sf21)* were found to have a muscle disorganization score of 0, and *dyc-1, alp-1, tln-1*, and *lev-11* had a score of 1 that were well correlated with the high scores of muscle function. The reason why these muscle mutants were not structurally disorganized might be because the mutation was not strong enough to result in any structural disorganization in muscle. Alternatively, it could be due to the specific protein location that our antibodies stained while the mutation resulted in muscle disorganization in another region that our protein of interest colocalized to. For example, here we used ATN-1 to visualize the depth of dense bodies of *zyx-1* mutants, yet it is reported that ZYX-1 proteins localize in the middle of dense bodies, and it might also be partially present in the basal region of the dense body where DEB-1 (vinculin) is located (60). Also, DYC-1 is known to localize at the edge of dense bodies and again DEB-1 might be a better candidate to visualize if there are any disorganizations in *dyc-1* animals (46). It was interesting to learn that despite the fact that DYC-1 localizes close to dense bodies, we only saw mild disorganization of M-lines of *dyc-1* mutants.

*unc-22(sf21)* was one of the mutants with completely normal muscle organization, as reported earlier (51). *sf21* is a missense mutation that inactivates the catalytic activity of the twitchin kinase domain, but expresses normal levels of the intact giant protein, twitchin. *unc-22(sf21)* mutants move faster, thus suggesting that the normal function of twitchin kinase activity is to inhibit muscle activity (51). Here, we saw that while they exert the same muscle force as WT as expected, the burrowing performance was mildly impaired. One possibility is that this impaired burrowing reflects a function of twitchin kinase in 3D but not 2D locomotion (e.g. force sensing of the environment). Interestingly, this muscle function and structural organization are in striking contrast to the other *unc-22* allele of *e66*,which was highly disorganized in sarcomere structure, is weak in muscle strength, and is severely impaired in burrowing. *e66* is a strong loss of function allele due to a 2 bp deletion resulting in a frame-shift and premature stop codon (51). It was previously suggested that twitchin plays both regulatory and structural roles in muscle, and our results add further evidence for this contention (13, 51).

*lev-11* encodes tropomyosin, and *lev-11(x12)* was another mutant with a very low muscle structural disorganization score of 1, slight burrowing impairment, and the same strength as WT. These results could also be expected due to the regulatory function of tropomyosin during actin-myosin interaction (61). This mutation resulted in no obvious structural disorganization; however, the burrowing assay has been the only locomotory assay so far that could detect such a regulatory impairment.

### *C. elegans* as a genetic model for muscle strength

The role of muscle strength in daily activities and exercise is indisputable. Several studies have investigated its importance in bone health (62) and its association with sarcopenia (63), heart diseases (64), and mortality (8). Age-related loss of muscle size and strength caused by a reduction in the size and number of individual muscle fibers (65) is associated with the increased frailty observed in older people (66). Given the economic impact of the aging population on healthcare systems (67), there is a greater need than ever to investigate the genetic basis of muscle strength.

Since the body wall muscle of *C. elegans* is functionally and structurally similar to human skeletal muscle and many of the muscle genes in *C. elegans* are known to have human homologs, our results highlight that *C. elegans* could be used as a model organism to study the genetics of muscle strength. Moreover, *C. elegans* is an established model for muscle aging (aka sarcopenia) (42, 68–70). Our physiological assays of burrowing and NemaFlex can effectively assess the muscle function in mutants with genetic defects in their muscle structure. By developing muscle function and muscle disorganization scores in *C. elegans*, we showed a strong correlation between the two suggesting that the lower muscle function score a mutant acquires, the higher level of structural disorganization it has. This result is significant since by utilizing forward and reverse genetic screens, both the muscle physiology and muscle structure can be evaluated in *C. elegans* to identify the genes responsible for muscle strength. We propose that muscle function score is a valuable means to pursue such investigations.

## Conclusions

Our results indicate the suitability of NemaFlex and burrowing assays to characterize muscle function of *C. elegans*. Using these testing approaches, we discuss the importance of the studied sarcomere proteins for muscle function and structure. We have shown that each of Pluronic gel burrowing assay and NemaFlex can report on different aspect of muscle physiology due to their 3D and 2D nature, stimulated and non-stimulated experimental conditions, with burrowing assay having a higher resolution in dissecting the muscle function defects. Interestingly, when these two assays are combined, they can inform more on the muscle function and structural organization. Thus the integrated scoring methodology we have developed enables further evaluation of *C. elegans* as a genetic model for muscle physiology to identify conserved genes responsible for human muscle strength.

## Declarations

### Ethics approval and consent to participate

Not applicable.

### Consent for publication

Not applicable.

### Availability of data and materials

All data generated or analyzed during this study are included in this published article and its supplementary information files. The video files and images used in this study are available from the corresponding author on reasonable request.

### Authors’ contributions

L.L. and K.K. performed NemaFlex and burrowing experiments. L.L. and H.Q. conducted immunostaining and captured images. L.L. and M.N. did statistical analysis and prepared figures. L.L., H.Q., C.M.R.L, N.J.S, G.M.B, and S.A.V discussed and interpreted the results. L.L. and S.A.V. wrote the paper. All authors read, edited and approved the final manuscript. L.L., C.M.R.L, G.M.B, N.J.S and S.A.V. conceptualized the project. S.A.V. supervised the study.

## Acknowledgements

We would like to thank Dr. Mathew Piasecki for useful discussions and Syed Tahsin Islam for assistance with data processing.

