## Supplementary information for "Investigating the correlation of muscle function tests and sarcomere organization in *C. elegans*"

Table S1. List of the mutants, their measured muscle strength, normalized muscle strength and body diameter

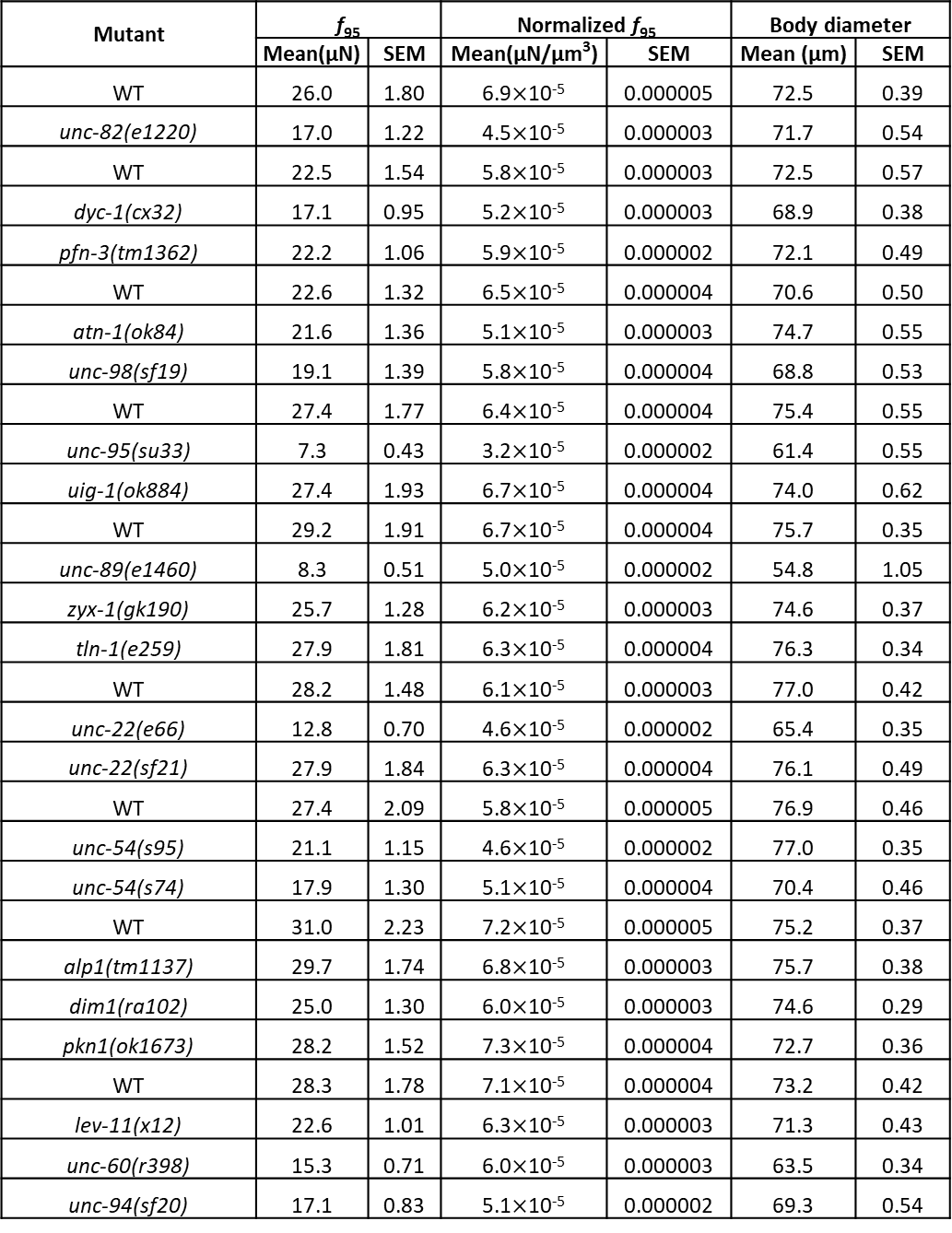

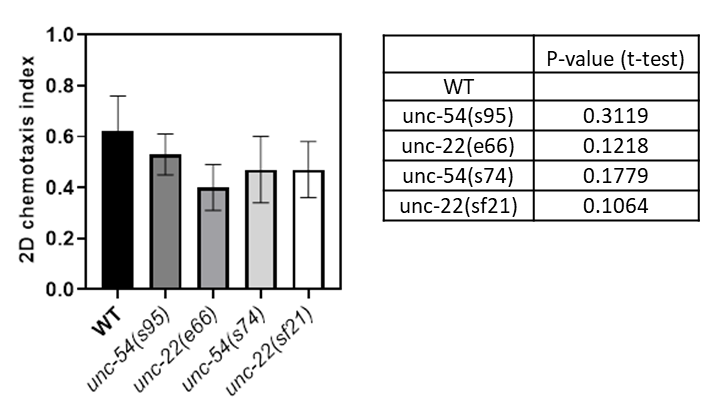

Figure S2. 2D chemotaxis indices of WT and Unc mutants. None of the Unc mutants were significantly different from WT in the standard 2D chemotaxis on agar plates. Statistical analysis was performed using a student’s t-test. Not significant: P-value>0.05

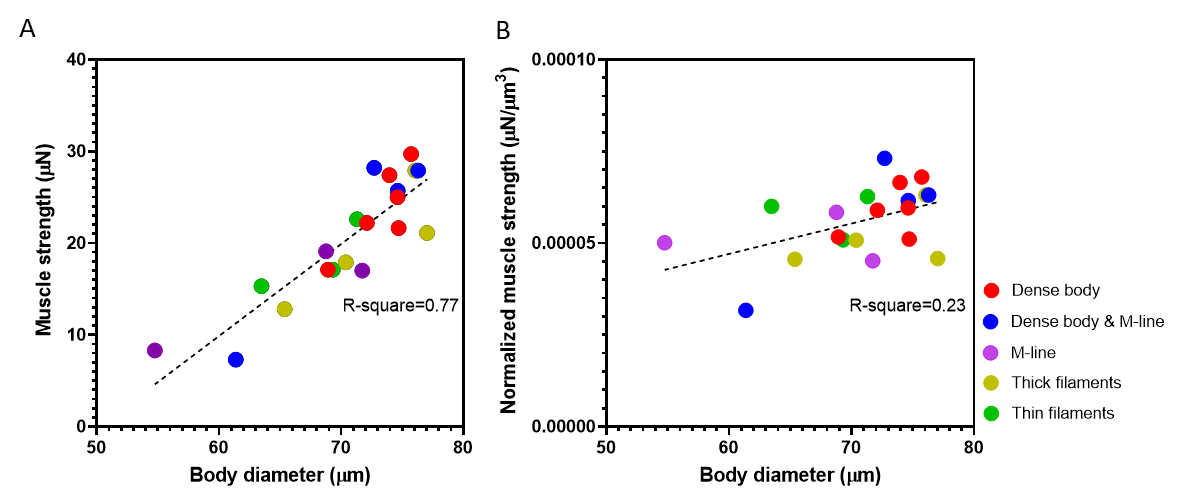

Figure S1. The effect of body diameter on (A) the measured muscle strength and (B) the normalized muscle strength which is measured muscle strength divided by the cube of body diameter.

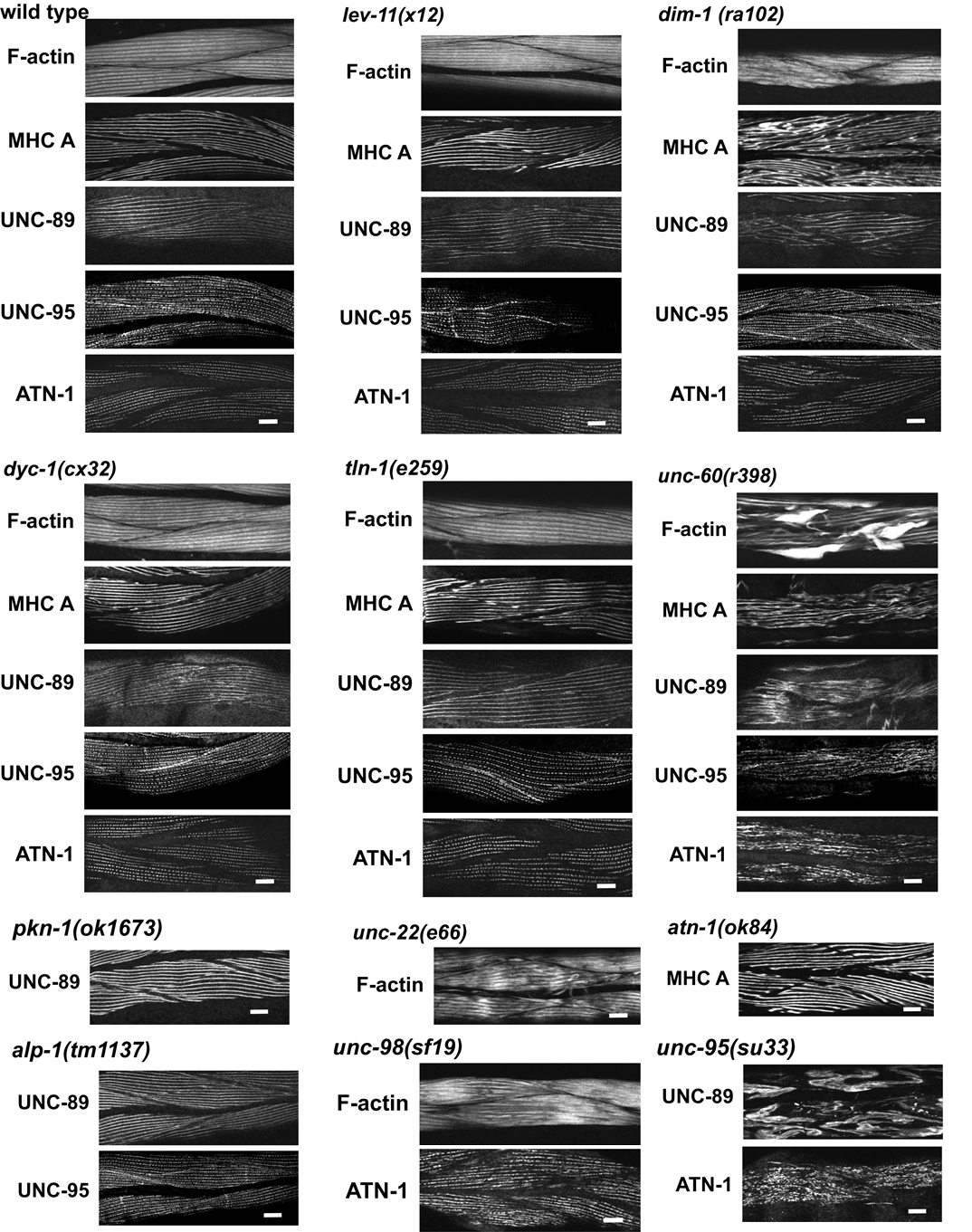

Figure S3. Phalloidin, MHCA, UNC-89, UNC-95, ATN-1 staining performed on sarcomere structural sites of wild type and muscle mutants .

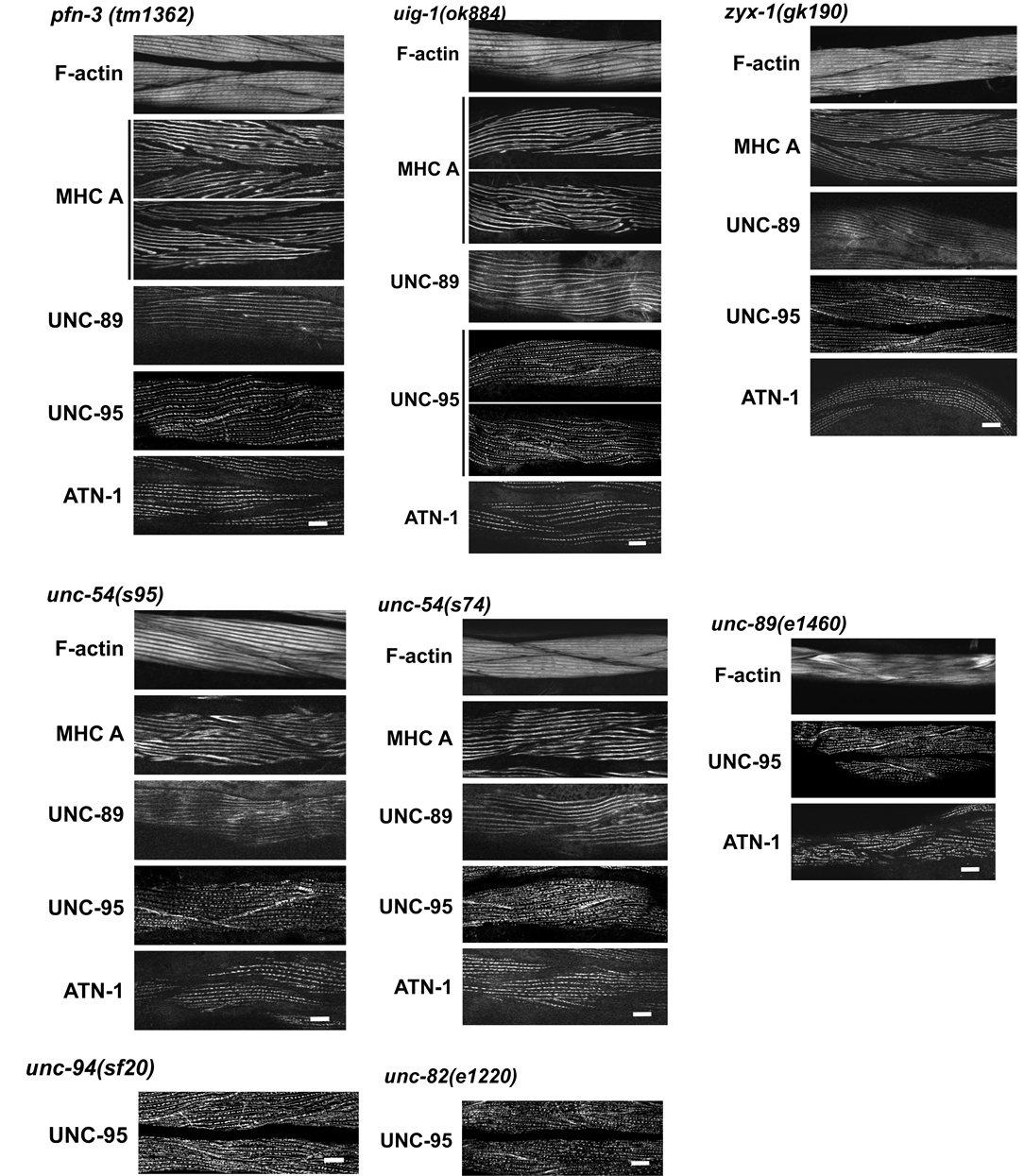

Figure S3-continued. Phalloidin, MHCA, UNC-89, UNC-95, ATN-1 staining performed on sarcomere structural sites of wild type and muscle mutants.

**Supplementary Note S1**

To define a muscle function score, we evaluated a linear combination of NemaFlex and burrowing Z-scores such that it explains most of the variations in the data. In order to find the best linear combination that preserve most of the variation in the data, principal component analysis (PCA) was used. First, NemaFlex and burrowing Z-scores were normalized to ensure that principal components are not affected by the magnitude of the measurements. To carry out the PCA, covariance matrix of normalized NemaFlex and burrowing Z-scores was calculated:

$C=\left[ \begin{matrix} 0.0813 & 0.0528 \\ 0.0528 & 0.0929 \end{matrix} \right]$

Where C is the covariance matrix of the data. Then, eigen-values and eigen-vectors of the covariance matrix were calculated:

$\lambda=\left[ \begin{matrix} 0.0339 & 0 \\ 0 & 0.1403 \end{matrix} \right] , v= \left[ \begin{matrix} -0.7449 & 0.6672 \\ 0.6672 & 0.7449 \end{matrix} \right]$

where columns of v include the eigen-vectors and the diagonal element in $\lambda$ contains the corresponding eigen-values. The amount of variance explained by each principal component is equal to the ratio of the corresponding eigen-value to sum of all eigenvalues. So, the principal component corresponding to eigen-value of 0.1403 contains 80 percent of the variation of the data (0.1403/(0.1403+0.0339)). Interestingly, this principal component suggests that the NemaFlex and burrowing data should be combined with almost equal weights (0.6672 and 0.7449). As a result, the muscle function score was simply defined as the sum of the individual normalized z-scores from the NemaFlex and burrowing assays.
